# Exchange protein directly activated by cAMP plays a critical role in regulation of vascular fibrinolysis

**DOI:** 10.1101/196899

**Authors:** Xi He, Aleksandra Drelich, Qing Chang, Dejun Gong, Yixuan Zhou, Yue Qu, Shangyi Yu, Yang Yuan, Jiao Qian, Yuan Qiu, Shao-Jun Tang, Angelo Gaitas, Thomas Ksiazek, Zhiyun Xu, Maki Wakamiya, Fanglin Lu, Bin Gong

**Affiliations:** Department of Pathology, University of Texas Medical Branch, Galveston, Texas 77555, USA; Department of Cardiovascular Surgery, Changhai Hospital, Shanghai 200433, China; Department of Mathematics and Statistics, Texas Tech University, Lubbock, Texas 79409, USA; Department of Neuroscience and Cell Biology, University of Texas Medical Branch, Galveston, Texas 77555, USA; Department of Electrical Engineering, Florida International University, Miami, Florida, 33199, USA. 14; Department of Neurology, University of Texas Medical Branch, Galveston, Texas 77555, USA

**Keywords:** EPAC1, vascular endothelial fibrinolysis, annexin A2, thrombosis, atomic force microscopy

## Abstract

**Rationale:** To maintain vascular patency, endothelial cells (ECs) actively regulate hemostasis. Among the myriad of pathways by which they control both fibrin formation and fibrinolysis is EC expression of annexin A2 (ANXA2) in a heterotetrameric complex with S100A10 [(ANXA2-S100A10)_2_]. This complex is a well-recognized endothelial surface platform for the activation of plasminogen by tissue plasminogen activator. A noteworthy advance in this field came about when it was shown that the cAMP pathway is linked to the regulation of (ANXA2-S100A10)_2_ in ECs.

**Objective:** These findings prompted us to determine whether a druggable target, namely the exchange <u>p</u>rotein directly <u>a</u>ctivated by <u>c</u>AMP (EPAC) pathway, plays a role in vascular luminal fibrinolysis.

**Methods and Results:** Taking advantage of our Epac1-null mouse model, we found that depletion of *Epac1* results in fibrin deposition, fibrinolytic dysfunction, and decreased endothelial surface ANXA2 in mice, which are similar to phenomena discovered in *ANXA2*-null and *S100A10*-null mice. We observed upregulation of EPAC1 and downregulation of fibrin in endocardial tissues beneath atrial mural thrombi in humans. Of note, our thrombosis model revealed that dysfunction of fibrinolysis in *EPAC1*-null mice can be ameliorated by recombinant ANXA2. Furthermore, we demonstrated that suppression of EPAC1 using a small-molecule inhibitor (ESI09) reduces the expression of ANXA2 in lipid rafts and impedes ANXA2 association with S100A10. Endothelial apical surface expression of both ANXA2 and S100A10 were markedly decreased in ESI09-treated ECs, which was corroborated by results from a nanoforce spectroscopy study. Moreover, inactivation of EPAC1 decreases tyrosine 23 phosphorylation of ANXA2 in the cell membrane compartment.

**Conclusions:** Our data reveal a novel role for EPAC1 in vascular fibrinolysis, by showing that EPAC1 is responsible for the translocation of ANXA2 to the EC surface. This process promotes conversion of plasminogen to plasmin, thereby enhancing local fibrinolytic activity.

## Introduction

Vascular endothelial cells (ECs) constitute the inner cellular lining of the vascular luminal wall, a highly dynamic and multi-functional interface between the inner vascular surface and flowing blood ^1-4^. To maintain vascular patency, ECs finely orchestrate the balance between blood coagulation and fibrinolysis on their luminal surfaces via complicated mechanisms, of which the plasmin-based fibrinolytic system has been well characterized ^5-8^. Fibrinolysis is elicited by the conversion of plasminogen to plasmin, which is activated by either of two plasminogen activators (PAs), tissue PA (tPA) or urokinase, on the surface of the fibrin thrombus or cell ^5, 6, 9^. Plasminogen is the pro-enzyme of the principal fibrinolytic protease plasmin, which is generated upon cleavage of plasminogen by PA at a single peptide bond at position Arg_560_ – Val_561_ ^10^ Besides secreting tPA on their surface, ECs express abundant plasminogen- and tPA-binding receptors ^7^, among which the annexin A2 (ANXA2) complex with S100A10 [(ANXA2-S100A10)_2_] is the best recognized and is emerging as the focus of research on a growing spectrum of biologic and pathologic processes ^9, 11-13^. On the endothelial luminal surface, (ANXA2-S100A10)_2_ recruits plasminogen and tPA, resulting in enhanced activation of plasminogen by at least 12-fold above baseline to produce fibrinolytic activity ^9, 11-13^. Moreover, *ANXA2*-null mice ^14^and *S10010A*-null mice ^8^ both demonstrate excessive accumulation of fibrin in the microvasculature.

ANXA2 is a Ca^2+^-regulated and phospholipid-binding protein that associates with biological membranes and the actin cytoskeleton ^11, 15, 16^. One of the well-recognized features of ANXA2 is its capacity to interact with its binding partner S100A10 and form the so-called heterotetrameric complex (ANXA2-S100A10)_2_, consisting of one S100A10 dimer and two ANXA2 molecules ^12, 17-19^. S100A10 is unique among the S100 protein family since it is locked in a permanent open conformation, which is comparable to the Ca^2+^-bound configuration of other members^11^. The ratio of monomeric ANXA2 to (ANXA2-S100A10)_2_ can vary among different cell types. In ECs, depletion of cellular ANXA2 results in rapid polyubiquitination of S100A10 for degradation in a proteasome ^17^. ANXA2 can associate with the cellular membrane by itself ^20^, yet, by binding S100A10, ANXA2 increases its sensitivity to Ca^2+^ and enhances its capability to bind the cellular membrane ^21^ and submembranous F-actin. ANXA2 and S100A10 are predominantly detected in cytoplasm and translocate across the cell membrane by an as yet unknown mechanism, which is independent from the classic endoplasmic reticulum—Golgi pathway ^22^. It has been proposed that ANXA2-containing multivesicular endosomes fuse directly with the cell membrane resulting in cell surface translocation ^23^. The process is regulated distinctively by phosphorylation of ANXA2 at different residues in its N-terminus in response to various stimuli ^22, 24, 25^. Therefore, the dynamics of ANXA2 in and near the plasma membrane compartment can govern vascular fibrinolysis on the endothelial surface in response to various signals ^11, 12, 21^. However, little is known about the precise mechanism mediating the cross-talk between environmental signals and the regulation of the dynamics of ANXA2 in a cell.

Cyclic adenosine monophosphate (cAMP) is an important molecular switch that translates environmental signals into regulatory effects in a cell ^26-30^. In multicellular eukaryotic organisms, the effects of cAMP are transduced by two ubiquitously-expressed intracellular cAMP receptors, the classic cAMP-dependent protein kinase A (PKA), and the more recently discovered exchange protein directly activated by cAMP (EPAC) ^31, 32^. To date, two isoforms of EPAC, EPAC1 and EPAC2, have been identified in humans. They are encoded by independent genes and predominantly expressed in different cell types. Both EPAC isoforms function by responding to increased intracellular cAMP levels in a PKA-independent manner and act on the same immediate downstream effectors, the small G proteins Rap1 and Rap2 ^33-35^. Based on accumulating evidence ^36-38^, the cAMP-EPAC signaling axis has been linked to vascular endothelial physiology and pathophysiology, including endothelial barrier function ^1, 39-43^, endothelial response to inflammatory stimuli ^44-46^, angiogenesis ^47, 48^, leukocyte adhesion to and migration across the endothelium 49-51, and intra-endothelial pathogen infection ^52^. Indeed, pharmacological and molecular approaches using human umbilical vein endothelial cells (HUVECs) have shown that EPAC1 is the major isoform in the ECs, and that Rap activation by EPAC1, and not by EPAC2, contributes to the effects of cAMP-elevating hormones on endothelial barrier functions ^1, 43, 53^. Of note, in ECs it has been documented that cAMP and EPAC are involved in hemostasis by driving the expressions of tPA and von Willebrand factor (vWF) ^25, 54, 55^. Recent reports showed that, in primary human ECs, (ANXA2-S100A10)_2_ is involved in the forskolin-induced cAMP-dependent secretion of vWF ^25^. Moreover, the Epac-Rap1 pathway can regulate exocytosis of the exocytotic organelles called Weibel-Palade bodies, in which vWF is stored ^55^. These findings prompted us to determine whether the cAMP-EPAC signaling axis plays a role in balancing blood coagulation and fibrinolysis on the vascular interior wall surface.

In the present study, by taking advantage of *EPAC1*-null mice and an EPAC-specific inhibitor, we show that EPAC1 plays a novel important role in maintaining vascular patency. Using biochemical and biomechanical assays, we found that EPAC1 exerts its regulation on vascular fibrinolysis via modulating endothelial surface expression of ANXA2 and its association with S100A10.

## Materials and Methods

### Patients and samples

In four cases of rheumatic mitral stenosis with chronic atrial fibrillation, left atrial mural thrombi were seen in the left atrial appendages during open heart surgeries for mitral valve replacements under extracorporeal circulation support at Changhai Hospital, the Second Military Medical University (Shanghai, China). After removing the thrombus, a 5 x 5 mm^2^ piece of endocardial tissue directly underneath the thrombus in the left atrial appendage was harvested. Tissue samples were flash frozen in liquid nitrogen and homogenized for immunoblotting assays by pulverizer (Spectrum Laboratories, Rancho Dominiquez, CA) as described previously^56^. The biopsy incision was closed with a 5-0 polypropylene suture. Similar tissue samples from normal donor hearts were used as normal controls. Informed written consent was obtained from each patient prior to study enrollment. This study was approved by the Committee on Ethics of Changhai Hospital.

### Atomic force microscopy (AFM)

The biomechanical properties of ANXA2 and S100A10 at the cell surface were studied using an AFM system (Flex-AFM, Nanosurf AG, Liestal, Switzerland) that utilized relevant antibody-functionalized AFM probes. Colloidal cantilevers with a 5 µm polystyrene bead were used (SHOCON-G-PS, Applied NanoStructures, Mountain View, CA) to measure surface protein interaction forces^57, 58^. The cantilevers were functionalized by incubation with anti-ANXA2 mAb or anti-S100A10 mAb at 100 g/L in 0.1 mol/L NaHCO_3_ (pH 8.6) overnight at 4°C. Unbound proteins were rinsed off using PBS. The exposed surface of the bead was blocked by BSA (Sigma, St. Louis, MO) at 500 g/L in PBS ^58^. AFM imaging and measurements were generally taken within 1 h after blocking. The spring constant of the cantilever was calibrated using the Sader method in air^59^. The cantilever spring constant varied between 0.10-0.15 N/m. Force spectroscopy was done in static force mode operating on 25 µm^2^ areas on a living cell surface. The functionalized cantilever was manipulated into contact with the surface of a confluent monolayer of HUVECs. The maximum compression force was set to 150 pN. The contact time was kept constantly at 500 ms before the cantilever was retracted at a constant pulling speed of 1 µm/s to measure the force-extension curve. In order to evaluate the effect of nonspecific unbinding force, confluent cell monolayers were pretreated with ANXA2 or S100A10 antibodies at 25 g/L for 30 min to block ANXA2 or S100A10 on the cell surface before the AFM measurements. We scanned five cells per group, each with a different cantilever. Force-distance curves were analyzed using the open source software Atomic J^60^.

### Statistics

Statistical significance was determined using Student’s t test. Results were regarded as significant if 2-tailed P values were < 0.05. All data are expressed as mean ± standard error of the mean.

## Results

### 1. Absence of the *EPAC1* gene results in fibrin accumulation in the microvasculature

To ascertain the role of EPAC1 *in vivo*, we generated *EPAC1*-null mice that carry the *Rapgef3* (also known as *EPAC1*) allele lacking exons 3-6. We confirmed that the mutant allele is null by Western immunoblotting analysis (WB) (**Fig. S1**). Similar to published models ^61, 62^, our *EPAC1*-null mice are viable, fertile, and without overt abnormalities.

*In vitro* evidences suggested EPAC1 controls vascular endothelial (VE)-cadherin-mediated cell junction formation^39-41^. Given an *in vivo* study showing that deletion of *EPAC1* inhibits endothelial barrier baseline in skin and intestine, but not heart ^61^, we assessed vascular integrity in brain and lung in our *EPAC1*-null 5 model. Compared to wild-type mice, the Evans blue assay revealed no differences in the baseline of extravascular dye in brain and lung parenchyma in *EPAC1*-null mice; immunofluorescent microscopy (IF) analysis displayed similar structures of vascular tight or adherens junctions (**Fig. S2**).

Excessive accumulation of intra- and extravascular fibrin is a hallmark of enhanced blood coagulation or incomplete fibrinolytic function caused by plasminogen or plasminogen activator deficiency in mice ^8, 14^. We employed immunohistochemistry (IHC) to measure fibrin levels in *EPAC1*-null and wild-type mice. Increased fibrin signals were detected within brain, lung, and renal blood vessels in *EPAC1*-null mice, while comparable staining was not detected in similar tissues from wild-type mice (**Fig. 1** and **Fig. S3**). WB displayed increased level of fibrin within tissues of lung and kidney in *EPAC1*-null mice, compared to wild-type mice (**Fig. 1**). Interestingly, IHC studies on tissues from wild-type mice treated with a small molecule EPAC-specific inhibitor, ESI09 (10 mg/kg/day, given intraperitoneally for 5 days) ^63^, recapitulated IHC findings in the *EPAC1*-null mice. Increased fibrin immunostaining within tissues of multiple organs was evident (**Fig. 1**). A similar phenomenon was not observed in wild-type mice treated with vehicle only. These data suggest that EPAC1 deficiency causes fibrin accumulation in a subset of tissues.

**Fig. 1:**
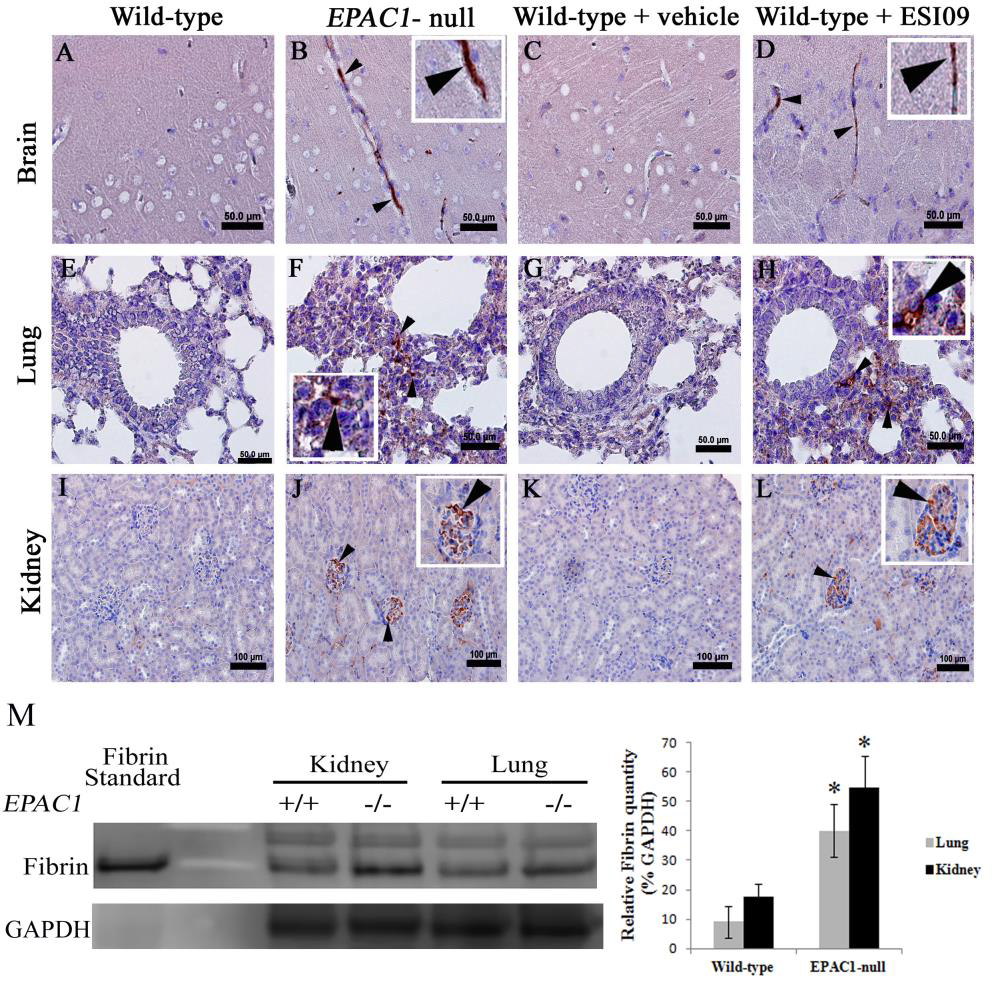
Absence of *EPAC1* gene or pharmacological inactivation of EPAC1 results in fibrin deposition in microvasculature in mice. Representative immunohistochemical analysis of brain (A-D), lung (E-H), and kidney (I-L) from wild-type and *EPAC1*-null mice, as well as wild-type mice treated intraperitoneally with ESI09 or vehicle only, is shown. Arrowheads indicate fibrin deposition in brain, lung, and renal blood vessels in both *EPAC1*-null and ESI09-treated wild-type mice. Normal mouse IgG was used as an antibody control (**Fig. S3**). Scale bars indicate 50 µm (A-H) or 100 µm (I-L). Western immunoblotting and densitometry analyses (M) of fibrin in mouse tissues from wild-type (n=4) and *EPAC1*-null (n=4) mice were normalized by GAPDH-specific control. *P < 0.05 compared with wild-type group.

We next examined blood coagulation function in *EPAC1*-null mice to determine whether fibrin accumulation is caused by enhanced blood coagulation. Prothrombin time (PT) and activated partial thromboplastin time (aPTT) are two biochemical indicators of blood coagulation ^8^. We observed that there were similar values of PT and aPTT between wild-type and *EPAC1*-null mice (**Fig. S4**), suggesting that *EPAC1* depletion does not affect the blood coagulation pathway characterized by these two assays. We also examined the platelet count and the platelet reactivity by measuring bleeding time^64^. We observed no differences between wild-type and *EPAC1*-null mice (**Fig. S4**).

Taken together, excessive accumulation of fibrin in the vasculatures in *EPAC1*-deficient mice is likely caused by impaired fibrinolytic activity rather than enhanced blood coagulation. Therefore, EPAC1 may play a regulatory role in baseline fibrinolytic homeostasis.

### 2. Reduced expression of EPAC1 in endocardial tissues underneath atrial mural thrombi in humans

Hypofibrinolysis was documented in rheumatic or non-rheumatic chronic atrial fibrillation patients, and associated with high thromboembolic risk ^65, 66^. We collected endocardial tissues (including cardiac endothelium) underneath surgery-confirmed left atrial mural thrombi from four cases of rheumatic mitral stenosis with chronic atrial fibrillation. The heart endocardium is primarily made up of cardiac ECs, which are derived from vascular ECs ^67^. Interestingly, WB analysis displayed that EPAC1 expression was prominently decreased, with increased levels of fibrin in the endocardial areas underneath the atrial mural thrombi compared to control tissue from normal donor hearts (**Fig. 2**). IF analysis displayed, compared to control tissue, enhanced fibrin deposition colocalized with reduced signal of EPAC1 in the endocardium and intima layer of coronary blood vessels in the cardiac wall (**Fig. 2**). This suggests that there is a correlation between reduced EPAC1 and deposition of fibrin in endothelium during formation of mural thrombi in human hearts.

**Fig. 2:**
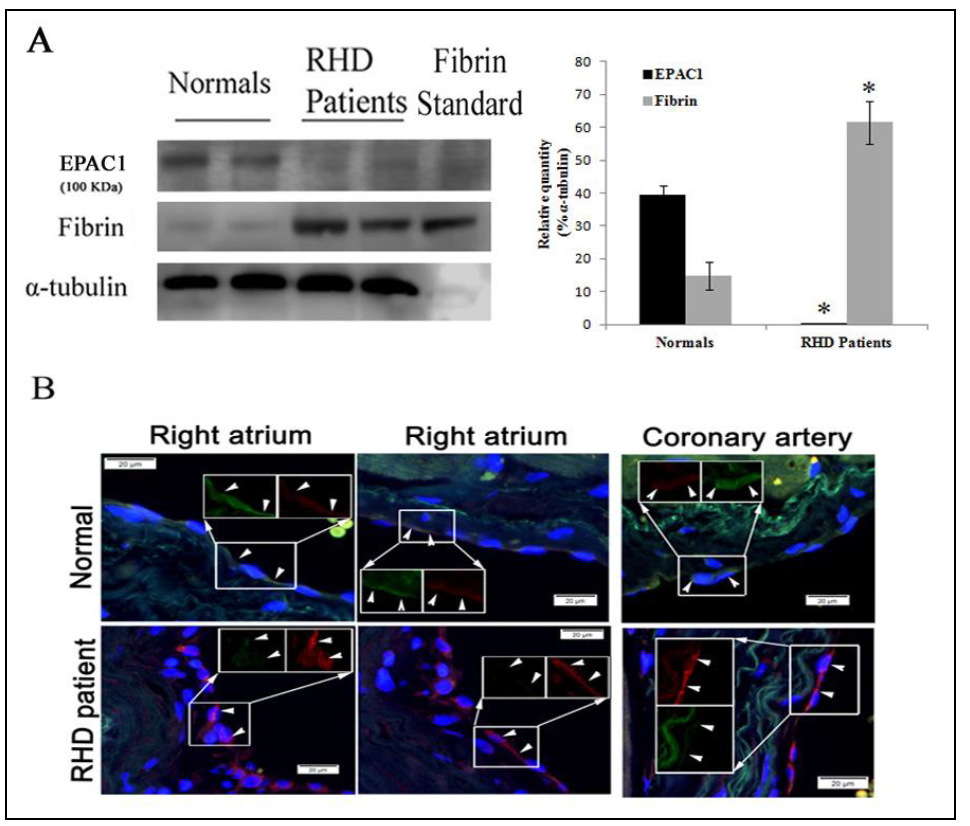
Reduced expression of EPAC1 and increased expression of fibrin in endocardial tissues underneath atrial mural thrombi in humans. Representative Western immunoblotting (A) shows markedly reduced expression of EPAC1 and increased expression of fibrin in endocardial areas (including cardiac endothelium) underneath atrial mural thrombi from patients with rheumatic heart disease (RHD) mitral stenosis with chronic atrial fibrillation compared to normal donor heart cases. Densitometry was used to quantify the relative intensity of EPAC1- and fibrin-specific immunoblots from patients (n=4) and normal cases (n=4) normalized by α-tubulin-specific controls. EPAC1 levels are presented as percentages of the indicated loading controls (*P < 0.01, compared to normal control). Representative dual-target IF staining localizes EPAC1 (green) and fibrin (red) in areas (arrowheads) of right atrial endocardium and intima layer of coronary blood vessel walls (B). Inserts depict split signals of EPAC1 (green) and fibrin (red) from the same area. Standard: human fibrin protein (Sigma-Aldrich). Scale bars indicate 20 µm

### 3. Deletion of *EPAC1* makes mouse susceptible to chemical-induced carotid arterial occlusion

The ferric chloride (FeCl_3_)-induced rodent carotid artery thrombosis model is a well-established experimental model for studying vascular fibrinolytic activity *in vivo*. In the model, the arterial occlusion caused by thrombus formation is measured by monitoring blood flow with a Doppler probe ^14, 68-70^. Using this model, we examined vascular fibrinolytic function in wild-type and *EPAC1*-null mice by comparing the maximal arterial occlusion (as % of baseline blood flow rate) (MaxO), maximal recovery of blood flow over 30 minutes after exposure to FeCl_3_ (as % of baseline blood flow rate) (MaxR) ^14^, and histological observation of FeCl_3_-induced carotid arterial thrombosis. Typical patterns of blood flow curves recorded during these experiments are illustrated (**Fig. 3**). Baseline blood flow was measured first, and there was no statistical difference between wild-type and *EPAC1*-null mice (1.9556 ± 0.1078 mL/min vs. 1.7694 ± 0.1265 mL/min, p=0.2748). Acute arterial thrombosis was induced by application of 7.5% FeCl_3_ to the adventitial arterial surface of the carotid artery^14^. Compared to wild-type mice, *EPAC1*-null mice demonstrated a significantly higher MaxO (36.40 ± 4.32 vs. 73.55 ± 7.95, P < 0.01) and a lower 6 MaxR (80.44 ± 4.97 vs. 31.49 ± 9.01, P < 0.01) for FeCl_3_-exposed carotid arteries. Furthermore, in a blinded study of hematoxylin and eosin (H&E)-stained cervical tissue, thrombi were detected in 8.33% of wild-type mice; in *EPAC1*-null mice, the incidence of detected thrombi was 0.42 (**Table**) (**Fig. S5**). These results indicate that *EPAC1*-null mice suffer more severe acute arterial occlusion caused by FeCl_3-_induced thrombosis.

**Fig. 3:**
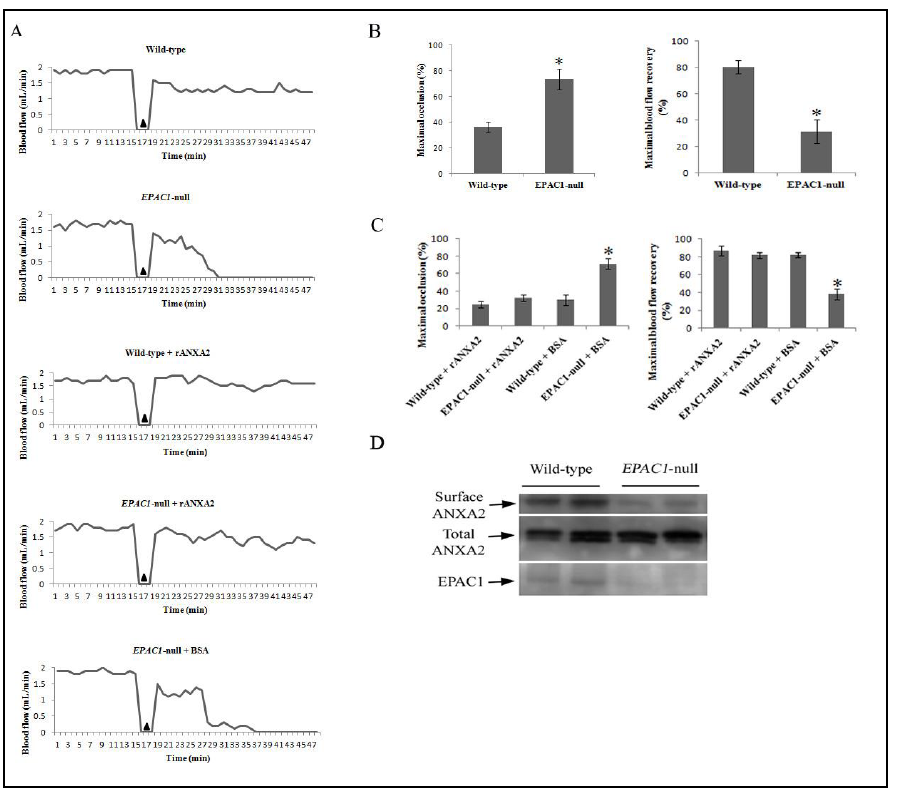
Impaired vascular fibrinolytic function can be restored by recombinant ANXA2 in *EPAC1* -null mice. Representative carotid blood flow before and after three-minute application of 7.5% FeCl_3_ (▴) (A). Compared to wild-type mice (n=12), *EPAC1*-null mice (n=12) demonstrated a significantly higher maximal occlusion (MaxO, *P < 0.01) and a lower maximal recovery (MaxR, *P < 0.01) (B). Wild-type (n=4) and *EPAC1-null* mice (n=11) were treated with rANXA2, showing no difference in MaxO and MaxR. Compared to *EPAC1*-null mice pretreated with BSA (n=5), *EPAC1*-null mice pretreated with rANXA2 show lower MaxO (*P < 0.01) and higher MaxR (*P < 0.01) (C). Western immunoblotting analysis showed reduced cell surface expression of ANXA2 *in vivo* (D)

The observation that *EPAC1*-null mice are susceptible to chemical-induced carotid arterial thrombosis, coupled with evidence that decreased expression of EPAC1 in human cardiac endothelia is associated with fibrin deposition on the surface, suggests that EPAC1 plays a role in regulation of vascular accumulation of fibrin.

### 4. Reintroduction of ANXA2 attenuates chemical-induced vascular occlusion in *EPAC1*-null mice

We next sought to delineate the cellular and molecular mechanisms by which EPAC1 might be involved in regulation of vascular accumulation of fibrin. EPAC1 is an intracellular sensor of cAMP. In our study, *EPAC1*-null mice showed an impaired ability to resist FeCl_3_-induced thrombosis, which is similar to the phenotypes observed in *ANXA2*-null mice ^14^ and *S100A10*-null mice ^8^. ANXA2 and S100A10 form a complex that is the most studied tPA binding receptor expressed abundantly in ECs ^7, 9, 11-13^. Furthermore, recombinant ANXA2 (rANXA2) can reduce thrombus formation in the FeCl_3_-induced rodent carotid artery thrombosis model ^68^. We introduced rANXA2 intravenously (i.v.) ^68^ into *EPAC1*-null mice to evaluate whether vascular occlusion can be restored.

Treatment with rANXA2 or bovine serum albumin (BSA) induced no changes in baseline blood flow in either wild-type or *EPAC1*-null mice (**Fig. 3**). After exposure to FeCl_3_, there was no significant difference in either MaxO or MaxR between wild-type and *EPAC1*-null mice when they were both pretreated with rANXA2 (**Fig. 3**). *EPAC1*-null mice pretreated with BSA showed decreased MaxO and increased MaxR of the FeCl_3_-exposed carotid artery, compared to the *EPAC1*-null mice pretreated with rANXA2. Blinded histological examination revealed that thrombi were detected in 40% of *EPAC1*-null mice pretreated with BSA; in *EPAC1*-null mice pretreated with rANXA2, the incidence of detected thrombi was 9.09% (**Table**). Moreover, we isolated biotinylated mouse aortic endothelial surface proteins from mice and revealed decreased cell surface ANXA2 in *EPAC1*-null mice (**Fig. 3**). These data indicate that attenuated blood flow resulting from *EPAC1* depletion correlates with impaired ANXA2-mediated endothelial surface fibrinolytic activity and can be restored by i.v. administration of rANXA2.

We also assessed the soluble part of the fibrinolytic system by measuring euglobulin clot lysis time ^71^ and found no differences between wild-type and *EPAC1*-null mice (**Fig. S4**). These observations suggest that the fluid-phase soluble fibrinolytic system was intact in *EPAC1*-null mice, a finding that is consistent with reduced extracellular ANXA2.

### 5. Inhibition of EPAC1 affects the interaction of ANXA2 with lipid rafts and impedes ANXA2 association with S100A10 in ECs

Given the fact that elevated levels of intra-endothelial cAMP can stabilize (ANXA2-S100A10)_2_ ^25^, we examined the correlation among EPAC1 expression and the levels and association of ANXA2 and S100A10. First, we performed qRT-PCR and immunoblotting to measure mRNA and protein levels of ANXA2 and S100A10 in tissues from wild-type and *EPAC1*-null mice. There was no significant difference in either mRNA or protein levels between wild-type and *EPAC1*-null mice in brain, lung, and kidney tissue (**Fig. S6**), suggesting that depletion of EPAC1 does not affect *de novo* protein synthesis of ANXA2 and S100A10 in these tissues.

Taking advantage of the EPAC-specific inhibitor ESI09, we determined the effect of EPAC1 inhibition on endothelial expression of ANXA2 and its partner S100A10 in the cellular membrane compartment. We treated HUVECs with ESI09 to inhibit EPAC1. Similar levels of mRNA and ANXA2 and S100A10 proteins were detected in vehicle- and ESI09-treated cells (**Fig. S6**), indicating no correlation between pharmacological inactivation of EPAC1 and *de novo* protein synthesis of ANXA2 and S100A10. However, immunoprecipitation assays with EC samples demonstrated that ESI09 treatment reduced associated ANXA2 in S100A10 precipitates, suggesting decreased formation of (ANXA2-S100A10)_2_ in ECs (**Fig. 4A**).

**Fig. 4:**
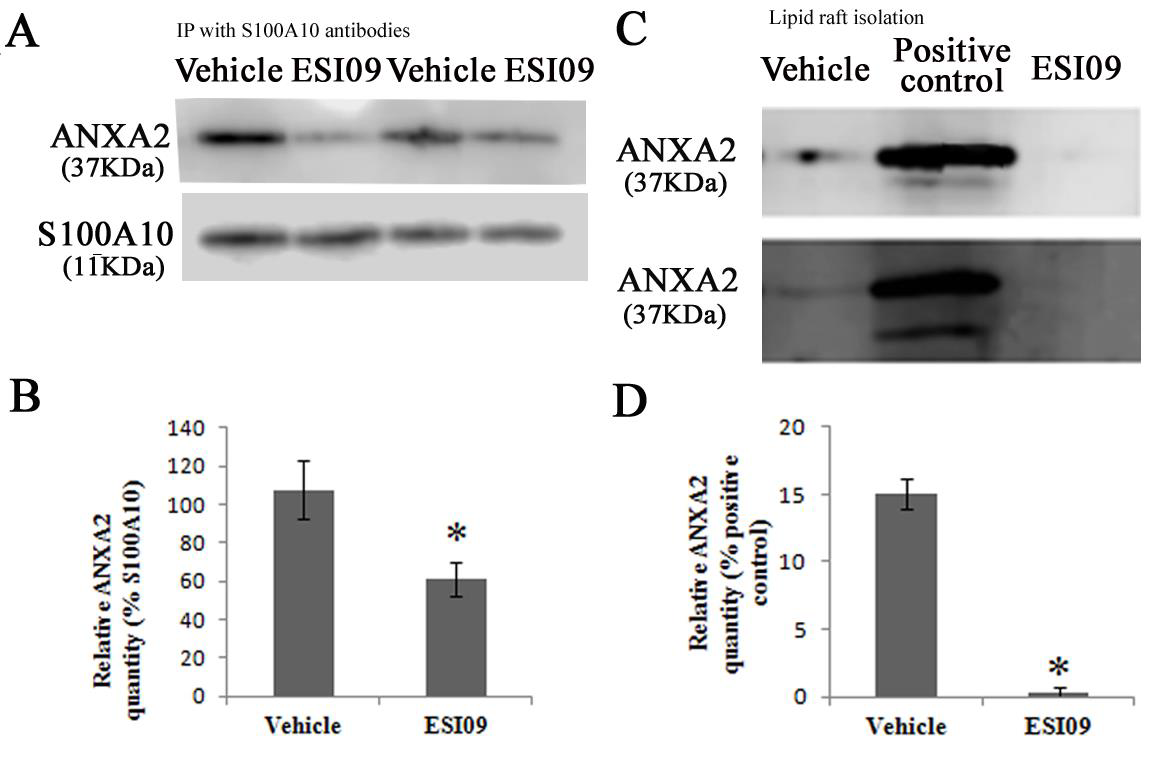
Inhibition of EPAC1 interrupts ANXA2 binding to lipid rafts and ANXA2 association with S100A10 in HUVECs. Western immunoblotting shows decreased levels of associated ANXA2 in S100A10 precipitates in ESI09-treated HUVECs, compared with vehicle-treated HUVECs (A) (*P < 0.05). ANXA2 levels were presented as percentages of the indicated loading controls in densitometry. Significantly reduced ANXA2 was detected in lipid rafts from ESI09-treated HUVECs, compared with vehicle-treated group (B) (*P < 0.01). A cell lysate was used as positive control. ANXA2 levels were presented as percentages of the positive controls in densitometry. Equal loading amounts of proteins were confirmed with lipid raft- and non-lipid raft-specific assays (**Fig. S7**). All experiments were repeated three times.

It has been established that ANXA2 binds to negatively charged phospholipids in cellular membranes. We therefore employed ultracentrifugation to isolate lipid rafts from membrane fraction samples of HUVECs in detergent conditions ^72^. We obtained highly purified lipid rafts (**Fig. S7**). In the vehicle-treated group, ANXA2 is still associated with lipid rafts after sample processing by ultracentrifugation. WB analysis revealed a significantly lower level of ANXA2 in lipid rafts in the ESI09-treated group compared to the vehicle-treated group (**Fig. 4B**).

These data suggest that inactivation of EPAC1 interrupts ANXA2 binding to the cell membrane and its association with S100A10.

### 6. Inhibition of EPAC1 decreases ANXA2 and S100A10 residing on EC apical surfaces

Both ANXA2 and S100A10 reside on the luminal surface of ECs mainly in the form of (ANXA2-S100A10)_2,_ a functional unit for plasmin-mediated fibrinolysis. To examine whether EPAC1 regulates the appearance of this crucial determinant for fibrinolytic activity on the EC luminal surface, we probed the expression levels of ANXA2 and S100A10 on the EC apical surface using biochemical and biomechanical approaches.

We performed impermeable cell-based ELISA assays ^73, 74^ to probe the cell surface levels of ANXA2 and S100A10. Endothelial surface levels of ANXA2 and S100A10 were determined as relative fluorescence units corrected for cell protein (FU/A550). Comparing vehicle- and ESI09-treated HUVECs in 96-well plates demonstrated that EPAC inhibition significantly decreased ANXA2- (P < 0.05) and S100A10-specific (P < 0.05) signals on the surfaces of HUVECs (**Fig. 5A**).

**Fig. 5:**
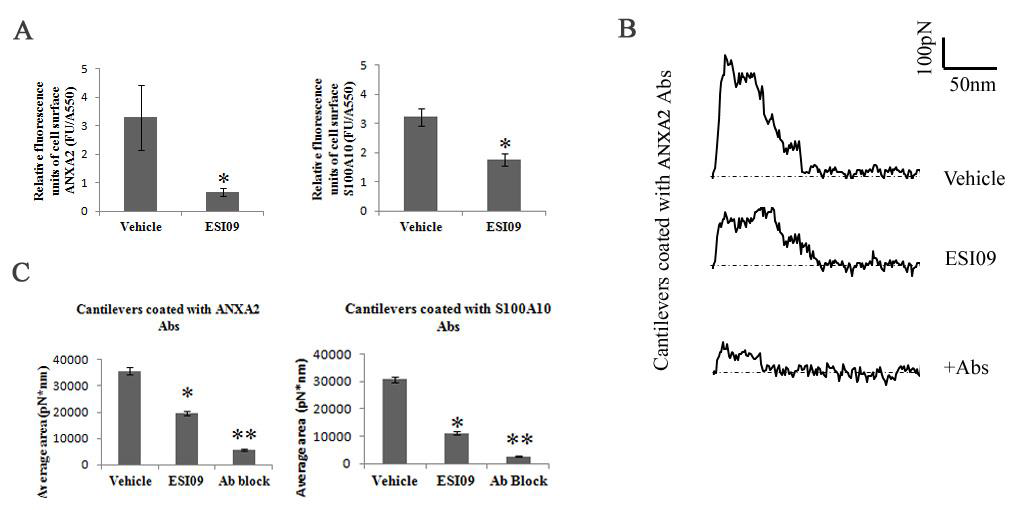
Inhibition of EPAC1 impedes ANXA2 and S100A10 residing on endothelial apical surfaces. An impermeable cell-based ELISA assay was employed to compare ANXA2 and S100A10 in ESI09- and vehicle-only treated HUVECs (A). Compared with the vehicle-treated group, ESI09-treated HUVECs show lower relative fluorescence units corrected for cell protein (FU/A550) of ANXA2 and S100A10 (* P < 0.05) signals on the surfaces of HUVECs. The data presented are representative of three independent experiments. The unbinding work of cell surface antigens with a relevant antibody on a cantilever surface was quantified by force-distance curve-based atomic force microscopy (AFM) (representative curves in B). During static force mode spectroscopy as the cantilever is pulled up from the cell surface, the unbinding force reaches its peak before all antigen–antibody interactions rupture. These specific unbinding force measurements can be compiled for quantification of total ANXA2 or S100A10 expression on selected areas on the HUVEC cell surface (C). With ANXA2 or S100A10 antibody-coated cantilevers, the unbinding work of vehicle-treated HUVECs is higher than that of ESI09-treated HUVECs (*P < 0.01), indicating reduced cell surface ANXA2 and S100A10 expression after ESI09 treatment. Ab block: ANXA2 antibody and S100D10 antibody at 25 g/mL for 30 min in medium, respectively, before the AFM measurement.

To further validate these results, we used atomic force microscopy (AFM) to quantify the distribution of ANXA2 and S100A10 on the surface of ECs. AFM is an advanced tool for studying biomechanical properties and has been used to determine the expression levels of cell surface proteins by measuring the binding affinity of specific protein–protein interactions, including antigen–antibody and receptor–ligand, with nanoforce spectroscopy **^58, 75^**. In our study, anti-ANXA2 or S100A10 antibodies were immobilized on polystyrene spheres attached to a colloidal cantilever. We measured the specific unbinding force during rupture of the interaction between the antigen (ANXA2 or S100A10) expressed at the apical surface of living HUVECs and the antibody-coated AFM cantilever probe. Interactions between antibodies on the AFM cantilever and cell surface antigens cause large adhesion forces, which are quantified by the deflection signal during separation of the cantilever from the cell. By tracking the cantilever deflection and retraction cycle, the binding, stretching, and ruptures of antibody–antigen complexes can be monitored in terms of the adhesive force changes on the cantilever over the distance traveled by the cantilever. Representative force-distance (FD) curves exhibit rupture events occurring during the interaction between antigens and antibodies in designated fields on the surface of a single living HUVEC (**Fig. 5B**). Therefore, by integrating the areas underneath the FD curve and above the baseline (zero force in **Fig. 5B**), we calculated the work that is required to break all interactive bonds between the cantilever and the EC, reflecting the quantity of antigen expression on the surface **^58, 75^**. Using an ANXA2 antibody-coated cantilever, greater unbinding work was measured on the surface of HUVECs exposed to vehicle compared with that of HUVECs exposed to ESI09 (3.568 ± 0.132 x 10^4^ pN*nm vs. 1.956 ± 0.072 x 10^4^pN*nm, P < 0.01), indicating that more ANXA2 antigens were detected on the surface of vehicle-treated HUVECs than ESI09-treated HUVECs (**Fig. 5C**). This suggests that EPAC1 inhibition reduces the amount of ANXA2 residing at the cell surface. Similarly, results obtained from using S100A10 antibody-coated cantilevers showed greater unbinding force on the surface of HUVECs exposed to vehicle compared to HUVECs exposed to ESI09 (3.075 ± 0.091 x 10^4^ pN*nm vs 1.119 ± 0.051 x 10^4^ pN*nm, P < 0.01), suggesting more S100A10 antigens were detected on HUVEC surfaces of the vehicle-treated group than the ESI09-treated group (**Fig. 5C**). The presence of the blocking antibody resulted in a significant decrease in the adhesion interaction between normal cells and the functionalized cantilever. These nano-biomechanical data corroborated the evidence gained from our biochemical studies that demonstrated that EPAC1 regulates the expression of ANXA2 and S100A10 on EC apical surfaces.

### 7. Inhibition of EPAC1 decreases tyrosine 23 phosphorylation of ANXA2 (Y23phANXA2) in the cell membrane compartment

ANXA2 binding to and translocation across the cell membrane are believed to be regulated via posttranslational modification of ANXA2, mainly phosphorylation or dephosphorylation of the N-terminal domain ^25, 76^. Phosphorylation of tyrosine 23 (Y23) influences ANXA2 binding to lipid rafts^23^. We examined Y23phANXA2 in HUVECs exposed to vehicle and ESI09. We observed that ESI09 decreased Y23phANXA2 in the membrane fractionation (**Fig. 6**), suggesting that EPAC1 regulates the dynamics of ANXA2, possibly by modulating phosphorylation or dephosphorylation of the N-terminal domain of ANXA2.

**Fig. 6:**
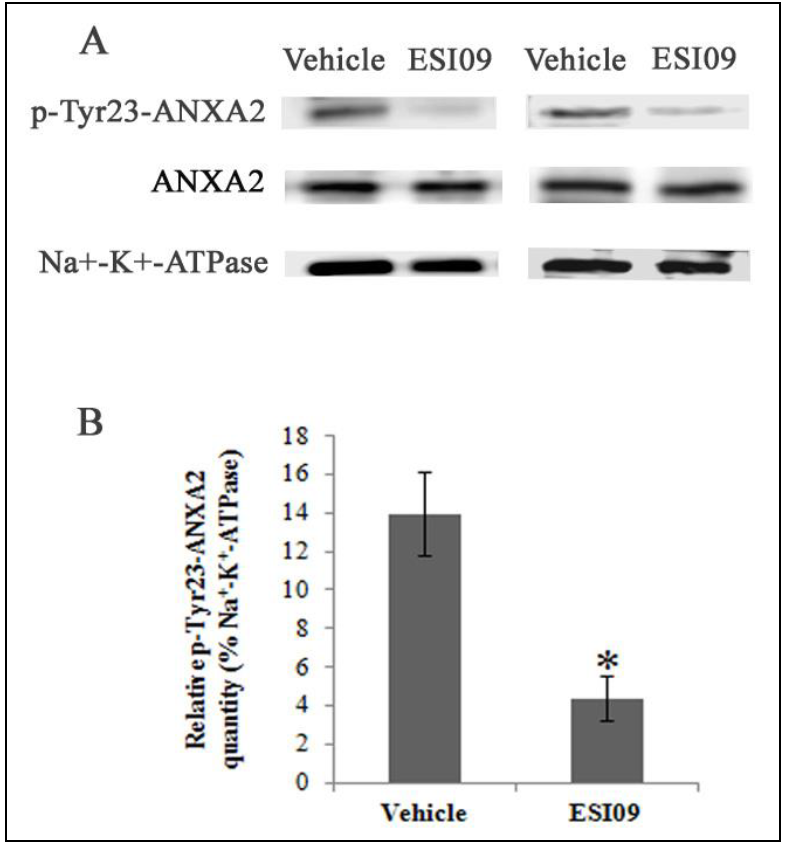
Inhibition of EPAC1 decreased tyrosine 23-phosphorylated ANXA2 in the cell membrane compartment. Representative Western immunoblotting (A) shows reduced expression of tyrosine 23-phosphorylated ANXA2 in the membrane fractionation of ESI09-treated HUVECs compared with vehicle-treated HUVECs (*P < 0.05). Densitometry was used to quantify the relative intensity of tyrosine23-phosphorylated ANXA2-specific immunoblots normalized by the indicated loading controls (B). All experiments were repeated three times.

## Discussion

Interrupted blood flow caused by a thrombus is the pathological foundation of some major human diseases. Fibrin, which is the main structural scaffold of a blood clot, is the ultimate product of coagulation and the major target of fibrinolysis^77^. Excessive accumulation of fibrin in tissue can be caused by enhanced blood coagulation or insufficiency in fibrinolytic function^8^. We detected increased fibrin accumulation in the microvasculature of multiple organs in *EPAC1*-null mice compared with wild-type mice, while there were no differences in measurements of PT and PTT assays, platelet count, and bleeding time. Reintroduction of ANXA2 attenuates chemical-induced vascular occlusion in *EPAC1*-null mice. This observation suggests that the increased fibrin accumulation detected in *EPAC1*-null mice is likely the result of impaired fibrinolysis rather than enhanced blood coagulation.

Plasmin, generated by cleavage of zymogen plasminogen, is the key protease involved in the dissolution of a fibrin clot. Among identified endothelial surface plasminogen and PA binding receptors, the (ANXA2-S100A10)_2_ has been potentially linked to the cAMP-EPAC signaling axis. In the context of coagulation-relevant regulation in HUVECs, the formation and stabilization of (ANXA2-S100A10)_2_ occurs in a cAMP-dependent manner^25^. The cAMP-EPAC pathway has been studied in cellular biological activities such as cell adhesion, cell junction, cell secretion, and cell differentiation^32, 38^. In ECs, EPAC1 and not EPAC2 is the dominant functional isoform^1, 43, 53^. EPAC1 is responsible for endothelial microtubule dynamics, cell secretion, angiogenesis, and barrier functions^1, 25, 40, 47^. In the present study, our *EPAC1*-null mice displayed excessive fibrin accumulation in the microvasculature of multiple organs. Compared with wild-type mice, *EPAC1*-null mice showed an inability to resist acute arterial occlusion caused by FeCl_3_-induced thrombosis. These results suggest that EPAC1 deficiency impaired fibrinolytic function. Our study, for the first time, showed the possible role of EPAC1 in regulating ECs fibrinolysis.

Interestingly, *ANXA2*-null mice ^14^ and *S100A10*-null mice ^8^ both also exhibited impaired fibrinolytic function, strongly suggesting that EPAC1 and (ANXA2-S100A10)_2_ may act cooperatively to mediate the fibrinolytic pathway. Employing biochemical and biomechanical technologies, we have obtained the following new lines of evidence: (1) inactivation of EPAC1 hampered the association between ANXA2 and S100A10 in ECs; (2) inactivation of EPAC1 decreased ANXA2 and S100A10 expression in ECs at the apical surface; and (3) AFM experiments revealed a dramatic reduction in the interactive forces between ANXA2-or S100A10-functionalized cantilevers and endothelial surfaces after inhibition of EPAC1 in ECs.

Taken together, these data reveal a tight association between EPAC1 and the dynamics of (ANXA2-S100A10)_2_ in ECs. (ANXA2-S100A10)_2_, which is expressed on both resting and activated ECs, possesses binding affinity for both plasminogen and tPA and serves as receptor for tPA-dependent plasmin generation on endothelial luminal surfaces ^9, 11-13^. Regulation of this heterotetramer, either by formation, stabilization, or translocation, modulates fibrinolytic activity on the EC luminal surface. The fact that *EPAC1*-null mice given rANXA2 intravenously gained the capability to attenuate acute arterial occlusion in chemical-induced thrombosis model further supports our interpretation that EPAC1 plays a critical role in (ANXA2-S100A10)_2_-mediated vascular fibrinolysis.

Despite our evidence supporting a connection between EPAC1 and (ANXA2-S100A10)_2_-mediated fibrinolysis, the underlying mechanism remains unclear. Translocation of (ANXA2-S100A10)_2_ occurs independently of the classical endoplasmic reticulum–Golgi pathway and does not involve *de novo* protein synthesis ^1, 22, 78^. The S100A10-binding motif is found in the N-terminal domain of ANXA2 ^12, 79^. ANXA2 lacks a typical signal peptide ^22^. ANXA2 in the cellular membrane compartment is believed to be regulated mainly via phosphorylation/dephosphorylation of the N-terminal domain of ANXA2 ^25, 76^,which is catalyzed by Ser/Thr kinases, Ser/Thr phosphatase, or Tyr kinase^22, 25, 76^. Brandherm *et al.* ^25^ identified that serine 11 dephosphorylation in ANXA2, catalyzed by a calcineurin-like phosphatase, is a positive switch for cAMP-dependent extracellular trafficking of (ANXA2-S100A10)_2_. Furthermore, He *et al*. reported that protein kinase C-activated phosphorylation of ANXA2 at serine 11 and 25 resulted in disassociation of the (ANXA2-S100A10)_2_ and prevented ANXA2’s translocation to the cell surface ^76^. Y23phANXA2 was identified as another regulatory switch during association of ANXA2 with lipid rafts^23^ and translocation of (ANXA2-S100A10)_2_^22^. After Y23phANXA2 through a Src-like tyrosine kinase-dependent pathway, the heterotetramer is translocated to the cell surface by an as yet unknown mechanism^22^. In this study we have shown that inactivation of EPAC1 by ESI09 decreased Y23phANXA2 in the EC membrane compartment, suggesting that EPAC1 may regulate ANXA2 phosphorylation/dephosphorylation and thus play a pivotal role in the dynamics of (ANXA2-S100A10)_2_. We are currently investigating the mechanism and pathway underlying this correlation. In conclusion, our studies combined an *in vitro* primary human endothelial system with an *in vivo* model and linked EPAC1 with the (ANXA2-S100A10)_2_-based fibrinolytic pathway to reveal a novel role for EPAC1 in vascular fibrinolysis. In ECs, EPAC1 is responsible for the formation and translocation of (ANXA2-S100A10)_2_, a key endothelial surface platform for tPA and plasminogen, aiding the conversion of plasminogen to plasmin.

## Acknowledgements

We gratefully acknowledge Dr. Edward Nelson for his important contributions for establishing the capacity of the atomic force microscopy (AFM) system, data analysis, and critical review of the manuscript. We gratefully acknowledge Drs. David Walker and Kimberly Schuenke for their critical reviews and editing of the manuscript. We thank Drs. Katherine Hajjar, Vladimir Motin, Vsevolod Popov, and Paul Boor for input during the planning phases of the experiments.

## Author contributions

B.G. and F.L. designed the study, performed experiments, analyzed data, and wrote the manuscript. F.L. and Z.X. contributed clinical samples, clinical correlative information, and performed *in vivo* experiments. X.H. and A.D. performed experiments, analyzed data, and wrote the initial draft of the manuscript. Q.C., D.G., J.Q., Y.Z., Y.Q., S.Y., Y.Y., and J.Q. performed experiments and analyzed data. Y.Q. and A.G. analyzed data. S.J.T. and T.K. assisted in designing experiments and provided comments and editorial changes. M.W. performed experiments, analyzed data, and provided comments and editorial changes.

## SUPPLEMENTARY MATERIALS

Materials and Methods

Fig. S1. Generation of EPAC1-null mice.

Fig. S2. Vascular integrity assessments.

Fig. S3. Normal mouse IgG was used as an antibody control during immunohistochemistry.

Fig. S4. Comparison of clotting parameters.

Fig. S5. Histological observation of FeCl_3_-induced carotid arterial thrombosis.

Fig. S6. Absence or inhibition of EPAC1 does not affect *de novo* protein synthesis of ANXA2 and S100A10.

Fig. S7. Lipid raft fraction-associated monosialotetrahexosyl ganglioside assay and non-lipid raft fraction-associated immunoblotting with anti-transferrin receptor.

---

**Table:** FeCl_3_-induced carotid arterial thrombosis detected in mice.

| Group | Pre-treatment | Number of animals | Carotid arterial thrombosis (%) * |
| --- | --- | --- | --- |
| Wild-type |  | 12 | 8.33 |
| *EPAC1-null* |  | 12 | 41.67 |
| Wild-type | BSA | 5 | 0 |
| *EPAC1-null* | BSA | 5 | 40 |
| Wild-type | rANXA2 | 4 | 0 |
| *EPAC1-null* | rANXA2 | 11 | 9.09 |

